# HIV-1 Nef synergizes with APOL1-G1 to induce nephrocyte cell death in a *Drosophila* model of HIV-related kidney diseases

**DOI:** 10.1101/2024.03.08.584069

**Authors:** Jun-yi Zhu, Yulong Fu, Joyce van de Leemput, Jing Yu, Jinliang Li, Patricio E. Ray, Zhe Han

## Abstract

**Background:** People carrying two *APOL1* risk alleles (RA) *G1* or *G2* are at greater risk of developing HIV-associated nephropathy (HIVAN). Studies in transgenic mice showed that the expression of HIV-1 genes in podocytes, and *nef* in particular, led to HIVAN. However, it remains unclear whether APOL1-RA and HIV-1 Nef interact to induce podocyte cell death.

**Method:** We generated transgenic (Tg) flies that express *APOL1-G1* (derived from a child with HIVAN) and HIV-1 *nef* specifically in the nephrocytes, the fly equivalent of mammalian podocytes, and assessed their individual and combined effects on the nephrocyte filtration structure and function.

**Results:** We found that HIV-1 Nef acts in synergy with APOL1-G1 resulting in nephrocyte structural and functional defects. Specifically, HIV-1 Nef itself can induce endoplasmic reticulum (ER) stress without affecting autophagy. Furthermore, Nef exacerbates the organelle acidification defects and autophagy reduction induced by APOL1-G1. The synergy between HIV-1 Nef and APOL1-G1 is built on their joint effects on elevating ER stress, triggering nephrocyte dysfunction and ultimately cell death.

**Conclusions:** Using a new *Drosophila* model of HIV-1-related kidney diseases, we identified ER stress as the converging point for the synergy between HIV-1 Nef and APOL1-G1 in inducing nephrocyte cell death. Given the high relevance between *Drosophila* nephrocytes and human podocytes, this finding suggests ER stress as a new therapeutic target for HIV-1 and APOL1-associated nephropathies.

**HIGHLIGHTS:**

- A new transgenic *Drosophila* model to study the pathogenesis of HIV-1-related kidney diseases with nephrocyte-specific expression of HIV-1 *nef* and an *APOL1-G1* risk allele derived from a patient with HIVAN.
- APOL1-G1 caused organelle acidification defects, reduced formation of autophagolysosomes, and reduced autophagy and protein aggregation, which culminated in ER stress.
- HIV-1 Nef induced ER stress through an autophagy-independent pathway. Furthermore, Nef and APOL1-G1 acted synergistically to heighten ER stress, which resulted in nephrocyte dysfunction and cell death.

**SIGNIFICANCE STATEMENT:** *APOL1* risk alleles are strongly linked to HIV-associated nephropathy (HIVAN) in people of African descent, but how HIV-1 and APOL1 interact and which pathways they might converge upon is unclear. A new *Drosophila* model to study HIV-1 Nef and APOL1-G1 (a risk allele) showed that Nef can induce ER stress in nephrocytes by itself, as well as exacerbate the organelle acidification defects and reduced autophagy induced by APOL1-G1, which further stimulates ER stress to a level that could cause nephrocyte cell death. Thus, we identified ER stress as the converging point for the synergy between APOL1-G1 and HIV-1 Nef in kidney cells, providing a potential therapeutic target for HIV-1 and APOL1-associated nephropathies.

## INTRODUCTION

HIV-associated nephropathy (HIVAN) is a kidney disease characterized by heavy proteinuria and rapid progression to chronic kidney failure ^1,2^. HIVAN renal histology shows collapsing glomerulopathy, de-differentiation and proliferation of podocytes and glomerular parietal epithelial cells, development of focal and segmental glomerulosclerosis (FSGS), and microcystic dilatation of renal tubules leading to kidney enlargement ^1,2^. People of African descent are at increased risk of developing HIVAN. Their risk to develop HIVAN or another chronic kidney disease is strongly associated with two *Apolipoprotein L1* risk alleles (*APOL1-RA*), *G1* and *G2* ^3,4^. People living with HIV-1 that carry two copies of the *APOL1-RA* have a ∼50% lifetime risk of developing HIVAN ^3–5^, and poorly controlled HIV-1 infection is the most powerful factor known to contribute to APOL1-kidney diseases ^4^. However, to date, the basic mechanisms through which the APOL1-RA interacts with HIV-1 to cause HIVAN remain unclear.

Mice and rats do not have an *APOL1* ortholog, nor can they be infected with HIV-1 ^6,7^. Nonetheless, transgenic (Tg) rodent models have been used to characterize the molecular mechanism through which the APOL1-RA and HIV-1 genes induce renal diseases ^8,9^. HIV-Tg26 mice and rats carrying a replication defective HIV-1 construct (driven by HIV-LTR lacking a 3kb sequence with the *gag* and *pol* genes, which are essential for viral replication), showed that the expression of HIV-genes in podocytes and tubular epithelial cells plays a critical role in inducing HIVAN in rodents ^10–15^. In addition, several HIV-Tg mouse models showed that the HIV accessory protein negative factor (Nef), is a key determinant of HIVAN pathogenesis ^10–15^. Studies in cultured kidney cells and *APOL1* Tg mice found that the expression levels and localization of APOL1 are crucial in determining its cytotoxicity ^17–20^. Notably, different dual APOL1-HIV-Tg26 mouse models have generated conflicting results, supporting either toxicity or protection ^17,21^. Thus, it remains unclear whether HIV-1 Nef and APOL1-RA interact directly to induce HIVAN, underlining the need for new animal models. *Drosophila* is an established model for kidney diseases and development ^18,22–27^. The fly nephrocyte is equivalent to the mammalian podocyte, with highly conserved filtration structure called the slit diaphragm ^28–31^. They also share many other similarities in endocytosis and exocytosis for the formation and maintenance of the filtration structure ^28–31^. A majority of genes associated with kidney diseases have functional homologs in flies that are required for nephrocyte function ^23,27^. The *Drosophila* nephrocyte has been well-established to study the pathogenesis of *APOL1*-associated nephropathies ^18,25,26,32,33^. Here, we generated Tg flies that express HIV-1 *nef* and *APOL1-G1*, the most frequent risk variant, specifically in nephrocytes. Our findings indicate that Nef and APOL1-G1 act synergistically to induce nephrocyte cell death through the ER stress pathway, providing a potential therapeutic target for HIV-1 and APOL1-associated nephropathies.

## METHODS

### Fly strains

Flies were maintained on standard food (Meidi LLC) at 22°C under standard conditions. The following *Drosophila* lines were obtained from the Bloomington Drosophila Stock Center (BDSC): *Dot*-Gal4 (ID_6903), UAS-YFP-*Rab5* (ID_24616), UAS-YFP-*Rab7* (ID_23270), UAS-mCherry-*Atg8a* (ID_37750), and *w*^1118^ (ID_3605). We previously generated the *Hand*-GFP flies labeling the nuclei of nephrocytes and cardiomyocytes ^34^.

The APOL1-G0 and APOL1-G1 (cDNA from a child with HIVAN) *Drosophila* lines were generated as described before ^18^. The HIV-1 Nef cDNA (pGM91) was obtained from the NIH AIDS Research and Reference Reagent Program (contributed by Dr. John Rossi) ^35^. To generate UAS-*nef* constructs, the *nef* allele cDNA was cloned into pUAST-attB and the transgene was introduced into a docking site by germline transformation. Flies expressing *Dot*-Gal4;UAS-*APOL1-*(*G0*/*G1*) were crossed with UAS-*nef* transgenic flies at 22°C.

### 10kD dextran and FITC-albumin uptake assay

Dextran and albumin uptake by nephrocytes was assessed *ex vivo* in (20-day) female adult flies to expose nephrocytes in artificial hemolymph containing 70 mmol/l NaCl, 5 mmol/l KCl, 1.5 mmol/l CaCl_2_, 4 mmol/l MgCl_2_, 10 mmol/l NaHCO_3,_ 5 mmol/l trehalose, 115 mmol/l sucrose, and 5 mmol/l HEPES (Sigma-Aldrich). Following a 20-minute incubation with 10kD Texas Red-dextran (0.05 mg/ml, Invitrogen) or FITC-albumin (100 mM, Sigma-Aldrich) and fixation (10 min) in 4% paraformaldehyde in phosphate buffered saline (4%PFA), the nephrocytes were mounted with VectaShield mounting medium (Vector Laboratories), and imaged.

### Nephrocyte size and number

Nephrocytes were dissected from 20-day-old female adult flies and kept in artificial hemolymph, followed by fixation (10 min) in 4%PFA, and imaged. Nephrocyte size was determined using the area measurement function in ImageJ ^36^ (version 1.52a). Nephrocyte numbers were manually counted using images of *Hand-GFP*.

### Immunochemistry

Female (20-day-old) flies were dissected and fixed [20 sec heat fix in 100°C artificial hemolymph for Polychaetoid (Pyd); or in 4%PFA for Ref(2)p, ubiquitin (FK2), and PDI]. Immunochemistry was carried out using established methods ^18^. Antibodies used in this study includes: mouse anti-Pyd antibody (1:100, Developmental Studies Hybridoma Bank); rabbit anti-Ref(2)P (1:100, ab178440, Abcam); mouse anti-ubiquitin FK2 (1:100, Enzo Life Sciences); rabbit anti-protein disulfide isomerase (PDI) (1:100, Sigma-Aldrich), and Alexa Fluor 555 (1:1,000, Thermo-Fisher-Scientific).

### LysoTracker assay

Fly nephrocytes were dissected in artificial hemolymph, incubated (20 min) with LysoTracker (Thermo-Fisher-Scientific), fixed (10 min) with 4%PFA, mounted with VectaShield, and imaged.

### TUNEL assay

Nephrocytes were dissected in artificial hemolymph and heat-fixed (100°C, 20 min) in 4%PFA, blocked with blocking buffer (50mM Tris-Cl pH 7.4, 0.1% Triton X-100 and 188mM NaCl), incubated with In Situ Cell Death detection kit (Roche), mounted with VectaShield and imaged.

### Confocal imaging

Confocal images were obtained using a ZEISS LSM900 microscope and ZEN Blue (edition 3.0) acquisition software. For quantitative comparison of intensities, common settings were chosen to avoid oversaturation and then applied across images for all samples within an assay. ImageJ ^36^ (version 1.52a) was used for all image processing.

### Statistical analysis

Statistical tests were performed using PAST.exe software (University of Oslo). Data were tested for normality using the Shapiro-Wilk test (α=0.05). Normally distributed data were analyzed by one-way ANOVA followed by Tukey-Kramer post-test for comparing multiple groups. Non-normal distributed data were analyzed by Kruskal-Wallis H-test followed by a Dunn’s test for comparisons between multiple groups. The results are presented as mean ± s.d. Statistical significance was determined as P<0.05. Details are provided in Supplementary File 1.

## RESULTS

### HIV-1 Nef exacerbates APOL1-G1-induced nephrocyte dysfunction

We previously generated transgenic *Drosophila* lines with nephrocyte-specific expression of *APOL1-G0* or the *APOL1-G1* risk allele derived from a child with HIVAN ^18^. Here, we generated transgenic flies with nephrocyte-specific expression of HIV-1 *nef* to explore how it alone or combined with *APOL1* affects the structure and function of nephrocytes.

We determined the capacity of dissected nephrocytes to filter and endocytose 10kD dextran or larger FITC-albumin particles. HIV-1 *nef* nor *APOL1-G0* showed a reduction in uptake (Figure 1A-C). Nephrocytes expressing *APOL1-G0*+*nef* or *APOL1-G1*+*nef* showed a significant decreased uptake of dextran and FITC-albumin compared to those expressing *nef, APOL1-G0*, or *APOL1-G1*; with the greatest reduction in *APOL1-G1*+*nef* nephrocytes (Figure 1A-C), reflecting the higher toxicity of the *APOL1-G1* risk allele. These results show that HIV-1 Nef exacerbates APOL1-induced nephrocyte dysfunction.

**Figure 1.**
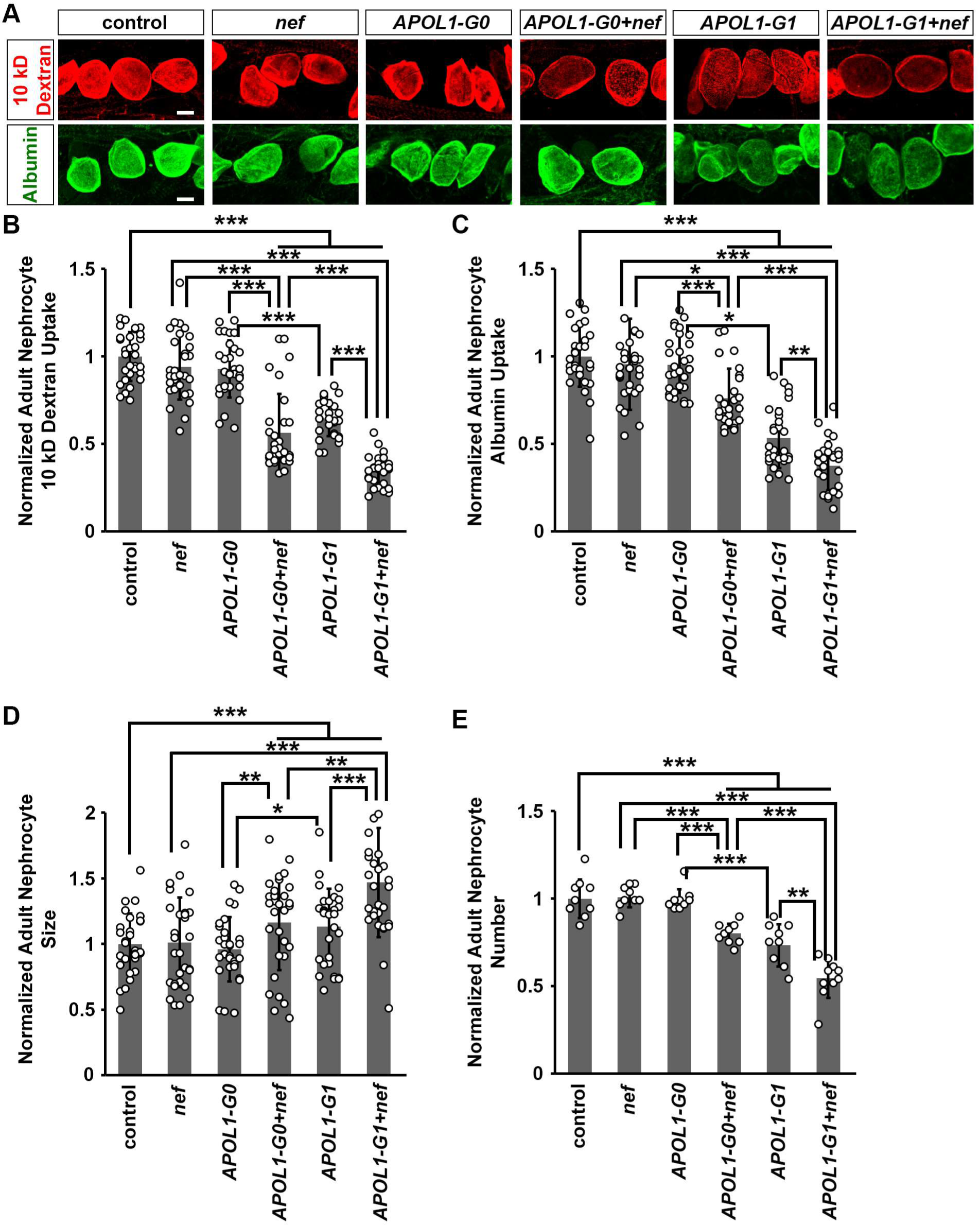
Expression of HIV-1 protein Nef facilitated nephrocyte function reduction, hypertrophy, and cell death due to APOL1-G1. **(A-E)** Flies used (20-day-old adult females): Control (*Dot*>*w*^1118^); *nef* (*Dot*>*nef*-OE); *APOL1-G0* (*Dot*>*APOL1-G0*-OE); *APOL1-G1* (*Dot*>*APOL1-G1*-OE); *APOL1-G0+nef* (*Dot*>*APOL1-G0*-OE+*nef*-OE); *APOL1-G1+nef* (*Dot*>*APOL1-G1*-OE+*nef*-OE). **(A) Top:** 10kD fluorescent dextran particle uptake (red) by nephrocytes using nephrocyte specific driver *Dot*-Gal4 to express HIV-1 *nef* alone or together with *APOL1-G0* and *APOL1-G1* at 22°C. Scale bar: 15 µm. **Bottom:** FITC-albumin particle uptake (green) by nephrocytes using nephrocyte specifically driver *Dot*-Gal4 to express HIV-1 *nef* alone or together with *APOL1-G0* and *APOL1-G1* at 22°C. Scale bar: 15 µm. **(B)** Quantitation of 10kD dextran uptake, relative to uptake in control flies. n=40 nephrocytes from 6 flies, per group. **(C)** Quantitation of FITC-albumin uptake, relative to uptake in control flies. n=40 nephrocytes from 6 flies, per group. **(D)** Quantitation of adult nephrocyte size, relative to size in control flies. n=40 nephrocytes from 6 flies, per group. **(E)** Quantitation of adult nephrocyte number, relative to cell number in control flies. n=10 flies, per group. Results have been presented as mean ± s.d., normalized to the control group. Kruskal–Wallis H-test followed by a Dunn’s test; statistical significance: *P<0.05, **P<0.01, ***P<0.001.

### Combined HIV-1 Nef and APOL1 cause nephrocyte hypertrophy and cell death

Previously, we showed that *APOL1-G1* induces progressive hypertrophy and accelerated cell death in nephrocytes ^18^. Hypertrophy validated here in 20-day-old fly nephrocytes expressing *APOL-G1* (Figure 1D). In contrast, nephrocytes expressing HIV-1 *nef* or *APOL1-G0*, did not show a significant change (Figure 1D), while those expressing *APOL1-G0*+*nef* or *APOL1-G1*+*nef* were significantly larger; with *APOL1G1*+*nef* nephrocytes the biggest (Figure 1D).

We observed reduced nephrocyte numbers in 20-day-old *APOL1-G1* Tg flies (Figure 1E). However, we did not observe significant changes in nephrocyte numbers in flies expressing HIV-1 *nef* or *APOL1-G0* (Figure 1E). In contrast, flies expressing *APOL1-G0*+*nef* or *APOL1-G1*+*nef*, showed a significant reduction in nephrocyte number compared to those expressing *APOL1-G0* or *APOL1-G1*; with numbers lowest in *APOL1-G1*+*nef* (Figure 1E). These results show that HIV-1 Nef combined with APOL1-G1 exacerbates the damage and loss of nephrocytes.

### HIV-1 Nef and APOL1-G1 combined disrupt the slit diaphragm filtration structure

The *Drosophila* slit diaphragm is a highly organized structure critical for the nephrocyte filtration function ^28^. Thus, we assessed the localization of the slit diaphragm protein polychaetoid (Pyd - the homolog of human tight junction protein ZO-1) ^30^. Control nephrocytes showed the typical fingerprint-like localization pattern of Pyd on the cell surface (Figure 2A), and the characteristic continuous circular ring in the optical medial view (Figure 2B). Notably, nephrocytes expressing *nef* or *APOL1-G0* did not show a visible disruption of Pyd (Figure 2A,B). However, nephrocytes expressing *APOL1-G0*+*nef* or *APOL1-G1*+*nef* showed highly disorganized Pyd localization, more so than either APOL-1 risk variant alone. Moreover, mislocalized Pyd was most evident in nephrocytes that expressed *APOL1-G1*+*nef* (Figure 2A,B), consistent with the altered uptake observed in nephrocytes expressing *nef* and *APOL1-G1* (Figure 1) which indicates a disrupted slit diaphragm filtration structure.

**Figure 2.**
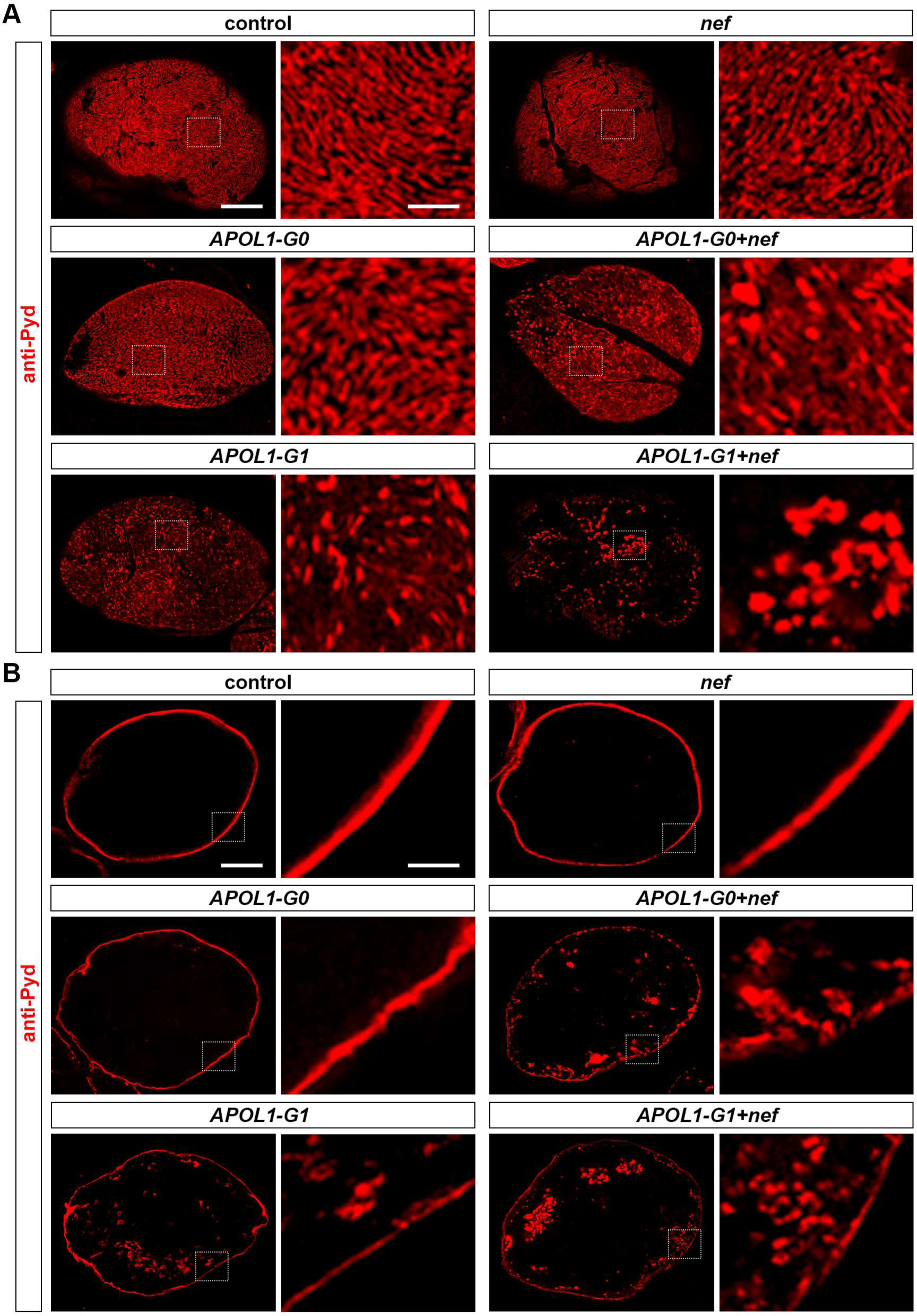
The expression of HIV-1 protein Nef in nephrocytes facilitated the disruption of the slit diaphragm due to APOL1-G1. **(A, B)** Flies used (20-day-old adult females): Control (*Dot*>*w*^1118^); *nef* (*Dot*>*nef*-OE); *APOL1-G0* (*Dot*>*APOL1-G0*-OE); *APOL1-G1* (*Dot*>*APOL1-G1*-OE); *APOL1-G0+nef* (*Dot*>*APOL1-G0*-OE+*nef*-OE); *APOL1-G1+nef* (*Dot*>*APOL1-G1*-OE+*nef*-OE). **(A, B)** Localization of slit diaphragm protein Polychaetoid (Pyd; red) at the surface (A) and medial (B) optical sections in nephrocytes using the nephrocyte specific driver *Dot*-Gal4 to express HIV-1 *nef* alone or together with *APOL1-G0* and *APOL1-G1* at 22°C. Boxed areas are shown magnified next to each image. Scale bar: 5 μm; Scale bar (magnified): 1 μm.

### Combined HIV-1 Nef and APOL1-G1 exert detrimental effects on nephrocyte endocytosis

We previously showed that HIV-1 can infect cultured human podocytes via dynamin-dependent endocytosis in a process facilitated by transmembrane TNF-α ^8^. Here, we assayed the expression of early (Rab5) and late (Rab7) endosomal markers in nephrocytes. Expression of *nef*, *APOL1-G0,* or *APOL1-G1*, nor combined, induced changes in the expression level or localization of Rab5 (Figure 3A,B). In contrast, Rab7 protein levels were significantly increased in nephrocytes expressing *APOL1-G1*, but not in those expressing *nef* or *APOL1-G0* (Figure 3C,D). Nephrocytes expressing *APOL1-G0*+*nef* or *APOL1-G1*+*nef* had significantly higher Rab7 protein levels; with the highest expression levels in *APOL1-G1*+*nef* nephrocytes (Figure 3C,D).

**Figure 3.**
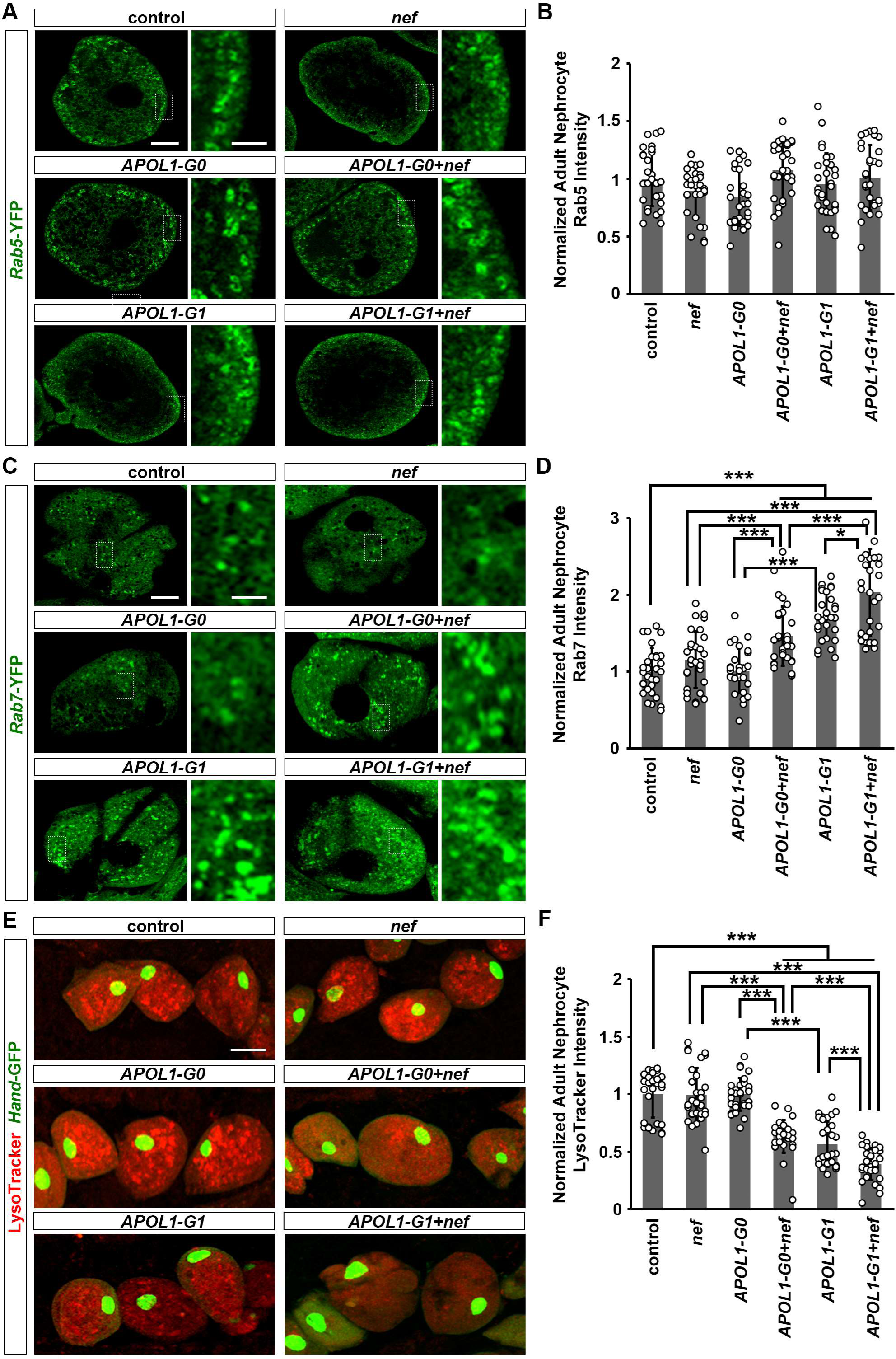
The expression of HIV-1 protein Nef in nephrocytes facilitated the disruption in endocytic membrane trafficking due to APOL1-G1. **(A, B)** Flies used (20-day-old adult females): Control (*Dot*>*Rab5*-YFP); *nef* (*Dot*>*Rab5*-YFP *+nef*-OE); *APOL1-G0* (*Dot*>*Rab5*-YFP*+APOL1-G0*-OE); *APOL1-G1* (*Dot*>*Rab5*-YFP*+APOL1-G1*-OE); *APOL1-G0+nef* (*Dot*>*Rab5*-YFP*+APOL1-G0*-OE+*nef*-OE); *APOL1-G1+nef* (*Dot*>*Rab5*-YFP*+APOL1-G1*-OE+*nef*-OE). **(C, D)** Flies used (20-day-old adult females): Control (*Dot*>*Rab7*-YFP); *nef* (*Dot*>*Rab7*-YFP *+nef*-OE); *APOL1-G0* (*Dot*>*Rab7*-YFP*+APOL1-G0*-OE); *APOL1-G1* (*Dot*>*Rab7*-YFP*+APOL1-G1*-OE); *APOL1-G0+nef* (*Dot*>*Rab7*-YFP*+APOL1-G0*-OE+*nef*-OE); *APOL1-G1+nef* (*Dot*>*Rab7*-YFP*+APOL1-G1*-OE+*nef*-OE). **(E, F)** Flies used (20-day-old adult females): Control (*Hand*-GFP, *Dot*>GFP+*w*^1118^); *nef* (*Hand*-GFP*, Dot*>*nef*-OE); *APOL1-G0* (*Hand*-GFP*, Dot*>*APOL1-G0*-OE); *APOL1-G1* (*Hand*-GFP*, Dot*>*APOL1-G1*-OE); *APOL1-G0+nef* (*Hand*-GFP, *Dot*>*APOL1-G0*-OE+*nef*-OE); *APOL1-G1+nef* (*Hand*-GFP*, Dot*>*APOL1-G1*-OE+*nef*-OE). **(A)** Expression of endocytosis-related proteins Rab5 (green, early endosome) in nephrocytes using the nephrocyte specific driver *Dot*-Gal4 to express HIV-1 *nef* alone or together with *APOL1-G0* and *APOL1-G1* at 22°C. Boxed areas are shown magnified next to each image. Scale bar: 5 μm; Scale bar (magnified): 1 μm. **(B)** Quantitation of Rab5 fluorescence intensity, relative to fluorescence in control nephrocytes. n=40 nephrocytes from 6 flies, per group. Note: None of the comparisons between groups were significant. **(C)** Expression of endocytosis-related proteins Rab7 (green, late endosome) in nephrocytes using the nephrocyte specific driver *Dot*-Gal4 to express HIV-1 *nef* alone or together with *APOL1-G0* and *APOL1-G1* at 22°C. Boxed areas are shown magnified next to each image. Scale bar: 5 μm; Scale bar (magnified): 1 μm. **(D)** Quantitation of Rab7 fluorescence intensity, relative to fluorescence in control nephrocytes. n=40 nephrocytes from 6 flies, per group. **(E)** LysoTracker dye fluorescence level (red) in nephrocytes using the nephrocyte specific driver *Dot*-Gal4 to express HIV-1 *nef* alone or together with *APOL1-G0* and *APOL1-G1* at 22°C. *Hand*-GFP transgene expression was visualized as green fluorescence concentrated in the nuclei of nephrocytes. Scale bar: 15 μm. **(F)** Quantitation of LysoTracker dye fluorescence intensity, relative to fluorescence in control nephrocytes. n=40 nephrocytes from 6 flies, per group. Results have been presented as mean ± s.d., normalized to the control group. Kruskal–Wallis H-test followed by a Dunn’s test; statistical significance: *P<0.05, ***P<0.001.

We also showed previously that *APOL1-G1* altered the acidification of organelles and the lysosomes in nephrocytes, which could impair their function ^18^. Here, we used LysoTracker to examine acidic vacuoles and lysosomes in nephrocytes with *nef* and *APOL1*. LysoTracker signal was significantly reduced in nephrocytes expressing *APOL1-G1*, but not *nef* or *APOL1-G0* (Figure 3E,F). Nephrocytes expressing *APOL1-G0*+*nef* or *APOL1-G1*+*nef* showed significantly lower LysoTracker signals (Figure 3E,F). Moreover, *APOL1-G1*+*nef* nephrocytes had the lowest signal (Figure 3E,F). These findings indicate that HIV-1 Nef exacerbates the changes induced by APOL1-G1 in key endocytic and acidification pathways that are essential for nephrocyte function.

### HIV-1 Nef enhances APOL1-G1-induced autophagic changes with accumulation of autophagosomes in fly nephrocytes

APOL1 can modulate autophagy in various experimental model systems ^17,19,25,32,37–39^. In *Drosophila*, autophagy-related 8a (Atg8a), the equivalent of human LC3, is a widely used marker for autophagy. Nephrocytes expressing *nef* showed no altered Atg8a-mCherry, but those expressing *nef* together with *APOL1* showed significantly higher levels of Atg8a-mCherry, with the highest signal in *APOL1-G1*+*nef* nephrocytes (Figure 4A,B). Likewise, we did not find altered refractory to Ref(2)P, a marker for cargo destined to be degraded by autophagy ^40^, in nephrocytes with *nef* alone (Figure 4C,D). In contrast, Ref(2)P was significantly increased in nephrocytes expressing *APOL1-G1*, and these changes were exacerbated in *APOL-G1*+*nef* nephrocytes (Figure 4C,D). In addition, we observed increased accumulation of ubiquitinylated proteins in nephrocytes expressing *APOL1-G1*, but not *nef* or *APOL1-G0* (Figure 4E,F). This accumulation was highest in *APOL1-G1*+*nef* nephrocytes (Figure 4E,F). These data demonstrate that HIV-1 Nef markedly increases APOL1-induced autophagy changes in nephrocytes.

**Figure 4.**
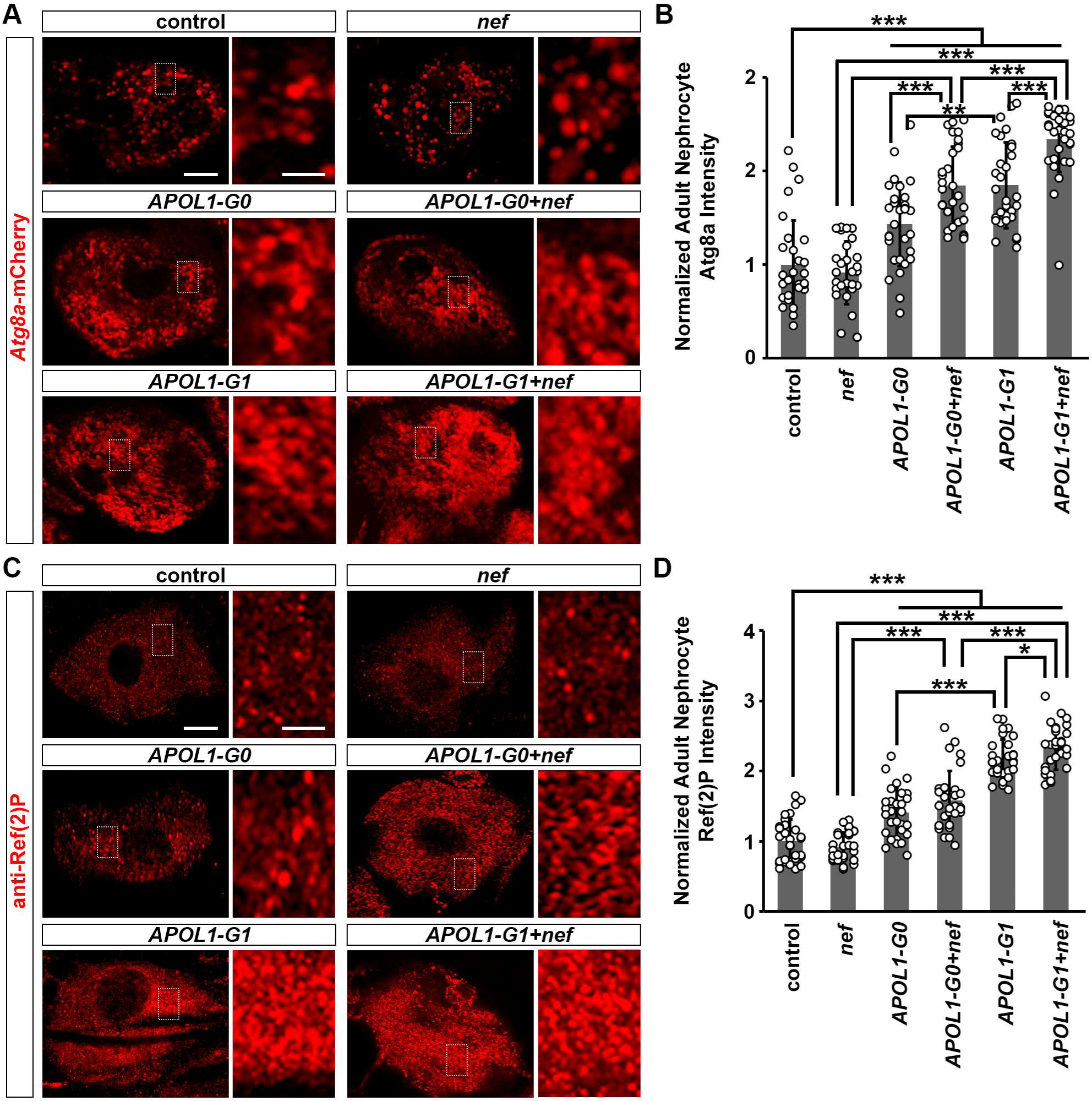
The expression of HIV-1 protein Nef in nephrocytes facilitated autophagy impairment due to APOL1-G1. **(A, B)** Flies used (20-day-old adult females): Control (*Dot*>*Atg8a*-mCherry); *nef* (*Dot*>*Atg8a*-mCherry*+nef*-OE); *APOL1-G0* (*Dot*>*Atg8a*-mCherry*+APOL1-G0*-OE); *APOL1-G1*(*Dot*> *Atg8a*-mCherry*+APOL1-G1*-OE); *APOL1-G0+nef* (*Dot*>*Atg8a*-mCherry*+APOL1-G0*-OE+*nef*-OE); *APOL1-G1+nef* (*Dot*>*Atg8a*-mCherry *+APOL1-G1*-OE+*nef*-OE). **(C-F)** Flies used (20-day-old adult females): Control (*Dot*>*w*^1118^); *nef* (*Dot*>*nef*-OE); *APOL1-G0* (*Dot*>*APOL1-G0*-OE); *APOL1-G1* (*Dot*>*APOL1-G1*-OE); *APOL1-G0+nef* (*Dot*>*APOL1-G0*-OE+*nef*-OE); *APOL1-G1+nef* (*Dot*>*APOL1-G1*-OE+*nef*-OE). **(A)** Expression of autophagy-related protein 8a (Atg8a; red, mCherry) in nephrocytes using the nephrocyte specific driver *Dot*-Gal4 to express HIV-1 *nef* alone or together with *APOL1-G0* and *APOL1-G1* at 22°C. Boxed areas are shown magnified next to each image. Scale bar: 5 μm; Scale bar (magnified): 1 μm. **(B)** Quantitation of Atg8a fluorescence intensity, relative to fluorescence in control nephrocytes. n=40 nephrocytes from 6 flies, per group. **(C)** Autophagy receptor refractory to sigma P (Ref(2)P; red) in nephrocytes using the nephrocyte specific driver *Dot*-Gal4 to express HIV-1 *nef* alone or together with *APOL1-G0* and *APOL1-G1* at 22°C. Boxed areas are shown magnified next to each image. Scale bar: 5 μm; Scale bar (magnified): 1 μm. **(D)** Quantitation of Ref(2)P fluorescence intensity, relative to fluorescence in control nephrocytes. n=40 nephrocytes from 6 flies, per group. **(E)** Expression of ubiquitinylated proteins (green) in nephrocytes using the nephrocyte specific driver *Dot*-Gal4 to express HIV-1 *nef* alone or together with *APOL1-G0* and *APOL1-G1* at 22°C. Boxed areas are shown magnified next to each image. Scale bar: 5 μm; Scale bar (magnified): 1 μm. **(D)** Quantitation of ubiquitinylated proteins fluorescence intensity, relative to fluorescence in control nephrocytes. n=40 nephrocytes from 6 flies, per group. Results have been presented as mean ± s.d., normalized to the control group. Kruskal–Wallis H-test followed by a Dunn’s test; statistical significance: *P<0.05, **P<0.01, ***P<0.001.

### HIV-1 Nef and APOL1-G1 induce ER stress in nephrocytes

APOL1-G1 can elicit the endoplasmic reticulum (ER) stress response ^26,32^, while HIV-1 Nef can bind the ER chaperone Calnexin (CNX) to induce ER stress ^41–43^. Therefore, we assessed protein disulfide-isomerase (PDI), a marker of ER stress in nephrocytes. PDI was significantly increased in nephrocytes expressing *nef* (Figure 5A,B). This increase was significantly more substantial when *nef* was expressed simultaneously with *APOL1-G1*; even when compared to *APOL1-G1* alone or *APOL1-G0*+*nef* nephrocytes (Figure 5A,B). Furthermore, nephrocytes that expressed *APOL1-G1*, but not *nef* or *APOL1-G0*, showed positive results in the TUNEL assay, a marker of cell death (Figure 5C). TUNEL-positive cells were also observed when nephrocytes expressed *APOL1-G0*+*nef* or *APOL1-G1*+*nef* (Figure 5C). This indicates that HIV-1 Nef alone can induce ER stress in *Drosophila* nephrocytes in an autophagy-independent manner, and that Nef can exacerbate the APOL1-G1-induced autophagy-dependent pathway to further stimulates ER stress, leading to nephrocyte dysfunction and cell death.

**Figure 5.**
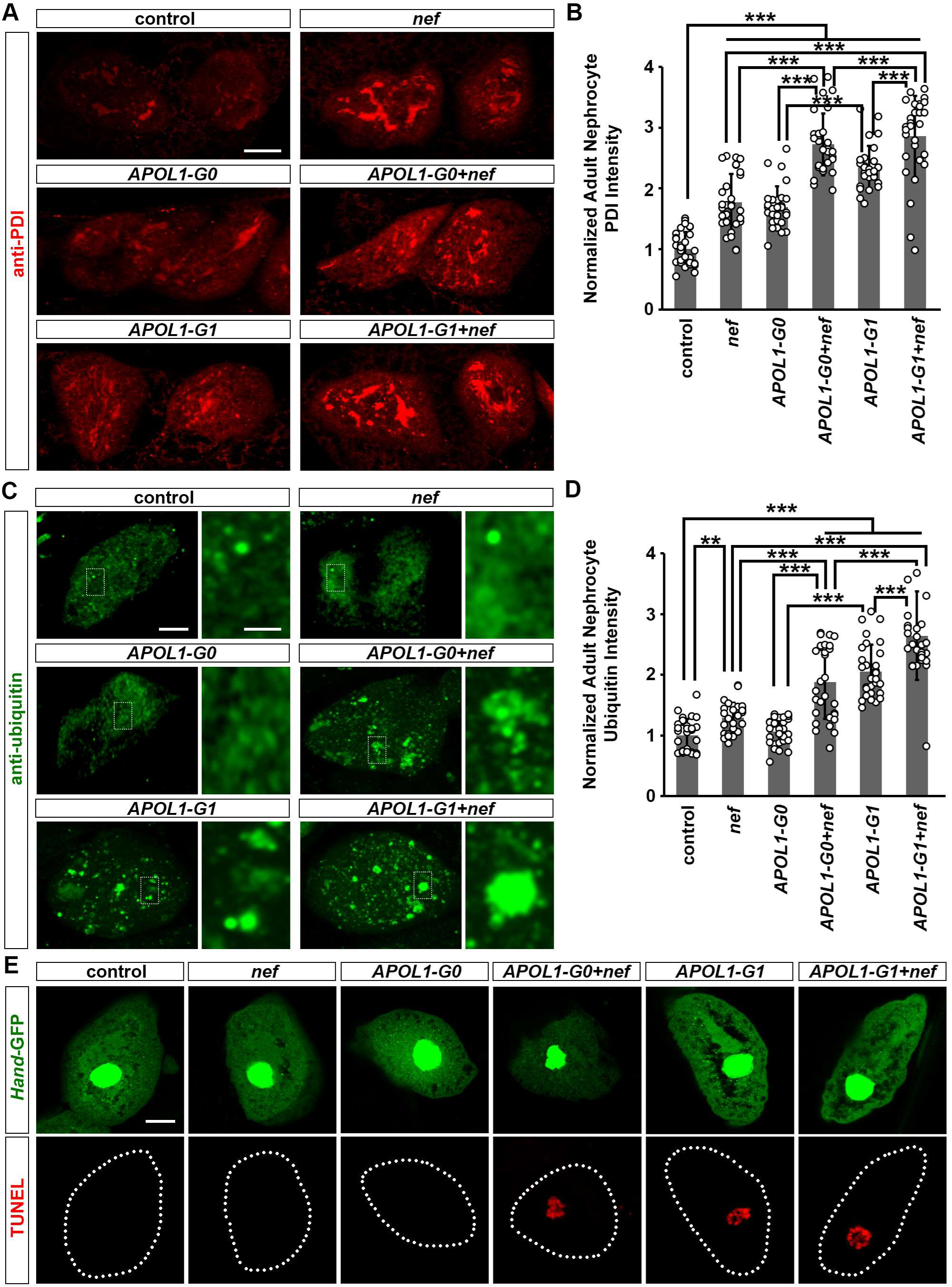
Expression of HIV-1 protein Nef in nephrocyte-induced ER stress. **(A, B)** Flies used (20-day-old adult females): Control (*Dot*>*w*^1118^); *nef* (*Dot*>*nef*-OE); *APOL1-G0* (*Dot*>*APOL1-G0*-OE); *APOL1-G1* (*Dot*>*APOL1-G1*-OE); *APOL1-G0+nef* (*Dot*>*APOL1-G0*-OE+*nef*-OE); *APOL1-G1+nef* (*Dot*>*APOL1-G1*-OE+*nef*-OE). **(C, D)** Flies used (20-day-old adult females): Control (*Hand*-GFP, *Dot*>GFP+*w*^1118^); *nef* (*Hand*-GFP*, Dot*>*nef*-OE); *APOL1-G0* (*Hand*-GFP*, Dot*>*APOL1-G0*-OE); *APOL1-G1* (*Hand*-GFP*, Dot*>*APOL1-G1*-OE); *APOL1-G0+nef* (*Hand*-GFP, *Dot*>*APOL1-G0*-OE+*nef*-OE); *APOL1-G1+nef* (*Hand*-GFP*, Dot*>*APOL1-G1*-OE+*nef*-OE). **(A)** ER stress marker protein disulfide isomerase (PDI; red) in nephrocytes using the nephrocyte specific driver *Dot*-Gal4 to express HIV-1 *nef* alone or together with *APOL1-G0* and *APOL1-G1* at 22°C. Scale bar: 15 μm. **(B)** Quantitation of PDI fluorescence intensity, relative to fluorescence in control nephrocytes. n=40 nephrocytes from 6 flies, per group. **(C)** Cell apoptosis marker TUNEL (red) in nephrocytes using the nephrocyte specific driver *Dot*-Gal4 to express HIV-1 *nef* alone or together with *APOL1-G0* and *APOL1-G1* at 22°C. *Hand*-GFP transgene expression was visualized as green fluorescence concentrated in the nuclei of nephrocytes. Scale bar: 5 μm.

**Figure 6.**
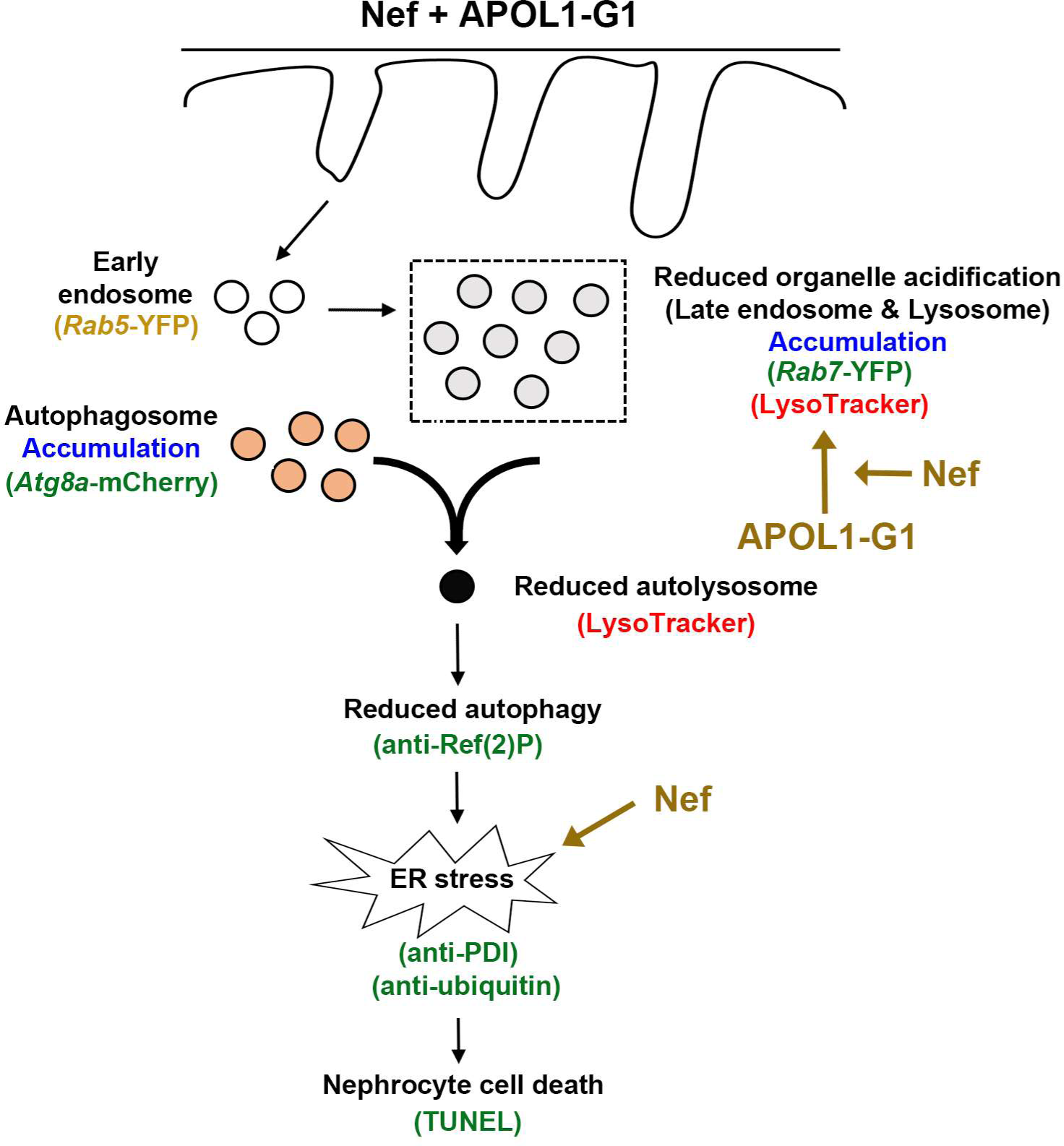
Model of HIV-1 Nef acting in synergy with APOL1-G1 through ER stress. Graphic depiction of the proposed model by which HIV-1 Nef induces ER stress, which in turn leads to cellular toxicity. Via a separate pathway, Nef exacerbates APOL1-G1-mediated ER stress. APOL1-G1 disrupts the late endosomes and lysosomes, which affects the fusion of autophagosomes and lysosomes, the reduced autophagy results in protein accumulation, leading to ER stress. Thus, when together HIV-1 Nef and APOL1-G1 significantly increase cellular toxicity through ER stress, which ultimately leads to cell death and tissue damage in the kidneys. (Green text indicates data showed upregulation, whereas red text indicates data showed downregulation, yellow text indicates no change.) APOL1-G1, Apolipoprotein L1-risk allele G1; Atg8a-mCherry, Autophagy-related 8a with mCherry fluorescent tag; ER, endoplasmic reticulum; PDI, protein disulfide isomerase; Rab5/7-YFP, Rab5 or Rab7 with yellow fluorescent protein tag; Ref(2)p, refractory to sigma P; TUNEL, terminal deoxynucleotidyl transferase dUTP nick end labeling assay.

## DISCUSSION

We have developed a new *Drosophila* system to explore how HIV-1 Nef and APOL1-G1 interact in nephrocytes. It revealed that HIV-1 Nef in addition to causing ER stress directly through an autophagy-independent pathway, also exacerbates the pathogenic effects of APOL1-G1 on the autophagy pathway, disrupting the function of nephrocytes and leading to their premature death.

APOL1-G1-induced cytotoxicity is not exclusively dependent on the nucleotide substitutions that define this risk variant. It is also affected by the BH3 domain ^26,44^, exon 4-encoded sequences ^37^, and different haplotypes ^45^. Therefore, we generated Tg flies expressing a common natural *APOL1-G1* derived from culture podocytes from a child with HIVAN ^14^. This APOL1-G1 contains the haplotypes (E150; I228; K255) and only differs at S342 and I384 from a common natural APOL1-G0 haplotype used to generate the control flies. Notably, this APOL1-G0 control haplotype is more toxic in cultured human kidney epithelial cells (HEK 293) than the reference APOL1-G0 haplotype ^14,19,25,46–52^. We previously showed that *APOL1-G0* expression in nephrocytes induced toxicity, albeit less severe compared to *APOL1-G1* ^18,32^. When we reduced the expression level of *APOL1-G0* in nephrocytes by lowering the temperature to 22°C, *APOL1-G0* toxicity was eliminated ^33^; hence all current fly experiments were conducted at 22°C (instead of 25°C used in our previous study) so that only *APOL1-G1* remains toxic. Nevertheless, we found that Nef could also synergize with APOL1-G0, making it toxic at 22°C (Figure 1), consistent with clinical studies showing that people of West African genetic ancestry living with a high HIV viral load can develop HIVAN even if they do not carry an *APOL1-RA* ^4^.

A study using *nephrin* promoter driven *APOL1-G0* Tg mice crossbred with HIV-Tg26 mice showed that fewer renal HIVAN lesions were developed compared to single HIV-Tg26 mice, suggesting a protective role for APOL1-G0 against HIV-induced nephropathy ^21^. However, studies in *APOL1* Tg mice with a BAC chromosome harboring *G0*, *G1,* or *G2* human *APOL1* found greater renal pathogenicity in *G2*/*G2* transgenic mice, compared to *G1*/*G1*; and that *APOL1-G0* did not rescue *APOL1-RA*-induced kidney injury when these mice were injected with Interferon-gamma (INF-γ) ^20^. Likewise, we found that *APOL1-G2* Tg flies developed more significant nephrocyte injury compared to *APOL1-G1* Tg flies ^32^. Overall, these studies indicate that the toxicity of APOL1-RA is mainly a function of its renal expression level ^20^, explaining why *APOL1* Tg mice driven by different promoters show different results ^9,17,20,53^. Our data is also consistent with a recent study using dual BAC/*APOL1*-HIV-Tg26 mice ^54^ to show that *APOL1-G0* did not minimize HIVAN lesions, whereas *APOL1-G1* aggravated them^54^. Previous studies showed that HIV-1 Nef can induce ER stress by binding a chaperon protein, Calnexin (CNX) ^41–43^. We showed that Nef caused ER stress in nephrocytes through an autophagy-independent pathway, likely by binding to a similar chaperon. In addition, we demonstrated that Nef together with APOL1-G1, disrupted the autophagy pathway (Figure 4) and promoting additional ER stress ^26,32^ (Figure 5). Defective endosomal trafficking has been reported in yeast and flies that expressed *APOL1-G1* ^25^. Likewise, immunofluorescence studies in cultured human podocytes revealed increased Rab7 (late endosomes) and LC3II (autophagosomes) positive organelles in cells expressing the high-risk (G1/G2) compared to low-risk (G0/G0) alleles ^17^. Studies in inducible podocyte-specific *APOL1-RA* transgenic mice also showed that G1 interferes with intracellular vesicular trafficking by impairing endocytic, autophagic, and acidification pathways, as well as by disrupting the maturation of autophagosomes and autophagic flux, which correlated with the development of FSGS ^17^. *APOL1-RA* was also shown to affect autophagy in cultured human podocytes ^65,66^.

Several studies have shown that HIV-1 can restrict autophagy to promote viral replication by inhibiting the fusion between autophagosomes and lysosomes ^38,55–57^. Nef overexpression in human astrocytes led to increased ATG8/LC3 ^56^, and impaired acidification in human cardiomyocytes, altering Rab7 localization and autophagy pathways ^57^. Alternatively, HIV-1 Nef reduced the fusion of autophagosomes to lysosomes by interacting with Beclin1 (BECN1)/Rab7 in cultured human cells ^58,59^. Together with our finding here, it appears that HIV-1 Nef-mediated inhibition of autophagic flux is conserved across cell types and species. Notably, the minor changes induced by Nef alone in nephrocytes resemble the mild podocyte changes observed in Tg mice with podocyte-specific *nef* expression ^12,60^. Whereas, other studies showed more severe podocyte injury when HIV-1 *nef* and *vpr* were expressed simultaneously in mouse podocytes^12^, suggesting that other HIV-1 genes and cytokines released by HIV infected cells are needed to evoke HIVAN phenotype fully. HIV-*nef* Tg mice driven by the CD4 promoter also developed kidney disease ^15,61,62^. Even though podocytes might not express HIV-1 genes ^15,61,62^, Nef can still be released by HIV-infected cells predominately in exosomes ^63^ and can be taken up by uninfected cells ^64^.

Our fly model has limitations, including that it is based on the ectopic overexpression of *APOL1* haplotypes, which can affect its intracellular localization and cytotoxicity ^7, 8^, and bypasses the infection or renal cells and the effects of other HIV-genes and circulating cytokines acting as second hits. HIVAN is a complex disease, involving many renal cell types and structures that cannot be studied in flies. However, our model permits the exploration of HIV-1 Nef and APOL1 interactions independently of many confounding variables. Most findings from Tg mice, cultured podocytes, and HIV-infected cells published so far are consistent with what we observed in nephrocytes. Since most relevant pathogenic pathways are highly conserved between fly nephrocytes and human podocytes, this new fly model provides a unique opportunity for rapid and cost-effective genetic or drug screens to better understand the APOL1 and HIV-1 interactions for inducing renal injuries, as well as to discover novel therapeutic treatment.

## ACKNOWLEDGMENTS

We thank the Bloomington Drosophila Stock Center (BDSC) based at Indiana University (Bloomington, IN) for the *Drosophila* stocks; and the Developmental Studies Hybridoma Bank (DSHB) based at the University of Iowa (Iowa City, IA) for providing the antibodies.

## AUTHOR CONTRIBUTIONS

J.Z. and Z.H. designed the study; J.Z., Y.F., J.Y. and JL. carried out the experiments; J.Z., J.vdL., and Z.H. analyzed and interpreted the data; J.Z. prepared the figures; J.Z, J.vdL., P.R., and Z.H. drafted and revised the manuscript; the manuscript has been critically reviewed and the final version approved by all authors.

