## Supplementary File 1 for "HIV-1 Nef synergizes with APOL1-G1 to induce nephrocyte cell death in a *Drosophila* model of HIV-related kidney diseases"

**Zhu et al.**
**HIV-1 Nef synergizes with APOL1 in cytotoxicity**
**Supplementary File 1**

### Fig. 1B

**1. Data normality using Shapiro-Wilk test**

| control | *nef* | *APOL1-G0* | *APOL1-G0+nef* | *APOL1-G1* | *APOL1-G1+nef* |
|---|---|---|---|---|---|
| 0.2276 | 0.6856 | 0.4701 | 0.0000416 | 0.6216 | 0.3275 |

**2. Nonnormal distributed data were analysed by Kruskal-Wallis H-test followed by a Dunn's test**

| | control | *nef* | *APOL1-G0* | *APOL1-G0+nef* | *APOL1-G1* | *APOL1-G1+nef* |
|---|---|---|---|---|---|---|
| control | | 0.1241 | 0.085 | 2.83E-08 | 1.09E-10 | 3.02E-11 |
| *nef* | | | 0.9117 | 3.52E-07 | 1.10E-08 | 3.02E-11 |
| *APOL1-G0* | | | | 4.11E-07 | 1.49E-08 | 3.02E-11 |
| *APOL1-G0+nef* | | | | | 0.001597 | 2.38E-07 |
| *APOL1-G1* | | | | | | 9.92E-11 |
| *APOL1-G1+nef* | | | | | | |

### Fig. 1C

**1. Data normality using Shapiro-Wilk test**

| control | *nef* | *APOL1-G0* | *APOL1-G0+nef* | *APOL1-G1* | *APOL1-G1+nef* |
|---|---|---|---|---|---|
| 0.5079 | 2E-05 | 0.1226 | 0.001348 | 0.01977 | 0.2759 |

**2. Nonnormal distributed data were analysed by Kruskal-Wallis H-test followed by a Dunn's test**

| | control | *nef* | *APOL1-G0* | *APOL1-G0+nef* | *APOL1-G1* | *APOL1-G1+nef* |
|---|---|---|---|---|---|---|
| control | | 0.3042 | 0.4464 | 8.29E-06 | 3.47E-10 | 4.50E-11 |
| *nef* | | | 0.7958 | 1.68E-04 | 1.86E-09 | 4.50E-11 |
| *APOL1-G0* | | | | 7.22E-06 | 7.38E-10 | 3.02E-11 |
| *APOL1-G0+nef* | | | | | 0.00001996 | 2.61E-10 |
| *APOL1-G1* | | | | | | 2.05E-03 |
| *APOL1-G1+nef* | | | | | | |

### Fig. 1D

**1. Data normality using Shapiro-Wilk test**

| control | *nef* | *APOL1-G0* | *APOL1-G0+nef* | *APOL1-G1* | *APOL1-G1+nef* |
|---|---|---|---|---|---|
| 0.9965 | 0.1226 | 0.4193 | 0.05167 | 0.06265 | 0.09574 |

**2. Nonnormal distributed data were analysed by Kruskal-Wallis H-test followed by a Dunn's test**

| | control | *nef* | *APOL1-G0* | *APOL1-G0+nef* | *APOL1-G1* | *APOL1-G1+nef* |
|---|---|---|---|---|---|---|
| control | | 0.9882 | 0.429 | 2.15E-02 | 6.35E-02 | 1.73E-07 |
| *nef* | | | 0.6952 | 7.48E-02 | 1.07E-01 | 1.53E-05 |
| *APOL1-G0* | | | | 7.62E-03 | 3.21E-02 | 8.35E-08 |
| *APOL1-G0+nef* | | | | | 0.3255 | 7.62E-03 |
| *APOL1-G1* | | | | | | 6.20E-04 |
| *APOL1-G1+nef* | | | | | | |

### Fig. 1E

**1. Data normality using Shapiro-Wilk test**

| control | *nef* | *APOL1-G0* | *APOL1-G0+nef* | *APOL1-G1* | *APOL1-G1+nef* |
|---|---|---|---|---|---|
| 0.5373 | 0.2992 | 0.0008726 | 0.9767 | 0.5244 | 0.1893 |

**2. Nonnormal distributed data were analysed by Kruskal-Wallis H-test followed by a Dunn's test**

| | control | *nef* | *APOL1-G0* | *APOL1-G0+nef* | *APOL1-G1* | *APOL1-G1+nef* |
|---|---|---|---|---|---|---|
| control | | 0.5403 | 0.8182 | 3.58E-04 | 3.58E-04 | 1.76E-04 |
| *nef* | | | 0.2444 | 1.95E-04 | 1.95E-04 | 1.70E-04 |
| *APOL1-G0* | | | | 1.71E-04 | 1.71E-04 | 1.72E-04 |
| *APOL1-G0+nef* | | | | | 0.2522 | 1.78E-04 |
| *APOL1-G1* | | | | | | 4.44E-03 |
| *APOL1-G1+nef* | | | | | | |

**Fig. 3B**

**1. Data normality using Shapiro-Wilk test**

| control | *nef* | *APOL1-G0* | *APOL1-G0+nef* | *APOL1-G1* | *APOL1-G1+nef* |
|---|---|---|---|---|---|
| 0.2555 | 0.0273 | 0.01537 | 0.1273 | 0.5407 | 0.08964 |

**2. Nonnormal distributed data were analysed by Kruskal-Wallis H-test followed by a Dunn's test**

| | control | *nef* | *APOL1-G0* | *APOL1-G0+nef* | *APOL1-G1* | *APOL1-G1+nef* |
|---|---|---|---|---|---|---|
| control | | 0.1624 | 0.01383 | 1.71E-01 | 4.29E-01 | 9.71E-01 |
| *nef* | | | 0.3328 | 5.08E-03 | 6.63E-01 | 1.49E-01 |
| *APOL1-G0* | | | | 7.70E-04 | 1.26E-01 | 1.27E-02 |
| *APOL1-G0+nef* | | | | | 0.04841 | 4.46E-01 |
| *APOL1-G1* | | | | | | 3.56E-01 |
| *APOL1-G1+nef* | | | | | | |

**Fig. 3D**

**1. Data normality using Shapiro-Wilk test**

| control | *nef* | *APOL1-G0* | *APOL1-G0+nef* | *APOL1-G1* | *APOL1-G1+nef* |
|---|---|---|---|---|---|
| 0.3656 | 0.3966 | 0.7306 | 0.01973 | 0.3053 | 0.0223 |

**2. Nonnormal distributed data were analysed by Kruskal-Wallis H-test followed by a Dunn's test**

| | control | *nef* | *APOL1-G0* | *APOL1-G0+nef* | *APOL1-G1* | *APOL1-G1+nef* |
|---|---|---|---|---|---|---|
| control | | 0.0993 | 0.7506 | 1.25E-05 | 2.03E-09 | 1.70E-09 |
| *nef* | | | 0.09626 | 5.08E-03 | 1.11E-06 | 5.09E-08 |
| *APOL1-G0* | | | | 3.32E-06 | 2.67E-09 | 6.12E-10 |
| *APOL1-G0+nef* | | | | | 0.006097 | 5.61E-05 |
| *APOL1-G1* | | | | | | 3.03E-02 |
| *APOL1-G1+nef* | | | | | | |

**Fig. 3F**

**1. Data normality using Shapiro-Wilk test**

| control | *nef* | *APOL1-G0* | *APOL1-G0+nef* | *APOL1-G1* | *APOL1-G1+nef* |
|---|---|---|---|---|---|
| 9E-05 | 0.055 | 0.8472 | 0.0003124 | 0.003982 | 0.6851 |

**2. Nonnormal distributed data were analysed by Kruskal-Wallis H-test followed by a Dunn's test**

| | control | *nef* | *APOL1-G0* | *APOL1-G0+nef* | *APOL1-G1* | *APOL1-G1+nef* |
|---|---|---|---|---|---|---|
| control | | 0.7731 | 0.3183 | 1.01E-08 | 6.01E-08 | 3.02E-11 |
| *nef* | | | 0.4825 | 5.97E-09 | 1.85E-08 | 5.49E-11 |
| *APOL1-G0* | | | | 1.61E-10 | 7.38E-10 | 3.02E-11 |
| *APOL1-G0+nef* | | | | | 0.1087 | 2.39E-08 |
| *APOL1-G1* | | | | | | 2.50E-03 |
| *APOL1-G1+nef* | | | | | | |

**Fig. 4B**

**1. Data normality using Shapiro-Wilk test**

| | control | *nef* | *APOL1-G0* | *APOL1-G0+nef* | *APOL1-G1* | *APOL1-G1+nef* |
|---|---|---|---|---|---|---|
| | 0.0021 | 0.0839 | 0.9456 | 0.03305 | 0.06512 | 0.0000655 |

**2. Nonnormal distributed data were analysed by Kruskal-Wallis H-test followed by a Dunn's test**

| | control | *nef* | *APOL1-G0* | *APOL1-G0+nef* | *APOL1-G1* | *APOL1-G1+nef* |
|---|---|---|---|---|---|---|
| control | | 0.865 | 0.0003563 | 7.09E-08 | 1.07E-07 | 2.61E-10 |
| *nef* | | | 0.00001868 | 3.82E-10 | 3.16E-10 | 1.09E-10 |
| *APOL1-G0* | | | | 1.77E-03 | 4.23E-03 | 4.57E-09 |
| *APOL1-G0+nef* | | | | | 0.9234 | 1.25E-05 |
| *APOL1-G1* | | | | | | 6.77E-05 |
| *APOL1-G1+nef* | | | | | | |

**Fig. 4D**

**1. Data normality using Shapiro-Wilk test**

| | control | *nef* | *APOL1-G0* | *APOL1-G0+nef* | *APOL1-G1* | *APOL1-G1+nef* |
|---|---|---|---|---|---|---|
| | 0.0406 | 0.0594 | 0.8527 | 0.0481 | 0.1123 | 0.7845 |

**2. Nonnormal distributed data were analysed by Kruskal-Wallis H-test followed by a Dunn's test**

| | control | *nef* | *APOL1-G0* | *APOL1-G0+nef* | *APOL1-G1* | *APOL1-G1+nef* |
|---|---|---|---|---|---|---|
| control | | 0.5493 | 0.00005265 | 4.44E-07 | 3.02E-11 | 3.02E-11 |
| *nef* | | | 2.028E-07 | 2.03E-09 | 3.02E-11 | 3.02E-11 |
| *APOL1-G0* | | | | 1.62E-01 | 1.29E-09 | 2.15E-10 |
| *APOL1-G0+nef* | | | | | 6.526E-07 | 3.35E-08 |
| *APOL1-G1* | | | | | | 3.39E-02 |
| *APOL1-G1+nef* | | | | | | |

**Fig. 4F**

**1. Data normality using Shapiro-Wilk test**

| | control | *nef* | *APOL1-G0* | *APOL1-G0+nef* | *APOL1-G1* | *APOL1-G1+nef* |
|---|---|---|---|---|---|---|
| | 0.006 | 0.2409 | 0.1805 | 0.007194 | 0.02723 | 0.0006126 |

**2. Nonnormal distributed data were analysed by Kruskal-Wallis H-test followed by a Dunn's test**

| | control | *nef* | *APOL1-G0* | *APOL1-G0+nef* | *APOL1-G1* | *APOL1-G1+nef* |
|---|---|---|---|---|---|---|
| control | | 0.002 | 0.2009 | 6.53E-08 | 6.07E-11 | 1.78E-10 |
| *nef* | | | 0.01501 | 1.53E-05 | 9.92E-11 | 5.57E-10 |
| *APOL1-G0* | | | | 1.60E-07 | 3.02E-11 | 3.82E-10 |
| *APOL1-G0+nef* | | | | | 0.234 | 5.71E-04 |
| *APOL1-G1* | | | | | | 1.04E-04 |
| *APOL1-G1+nef* | | | | | | |

**Fig. 5B**

**1. Data normality using Shapiro-Wilk test**

| | control | *nef* | *APOL1-G0* | *APOL1-G0+nef* | *APOL1-G1* | *APOL1-G1+nef* |
|---|---|---|---|---|---|---|
| | 0.0774 | 0.085 | 0.01637 | 0.02789 | 0.01487 | 0.001584 |

**2. Nonnormal distributed data were analysed by Kruskal-Wallis H-test followed by a Dunn's test**

| | control | *nef* | *APOL1-G0* | *APOL1-G0+nef* | *APOL1-G1* | *APOL1-G1+nef* |
|---|---|---|---|---|---|---|
| control | | 9E-09 | 1.695E-09 | 3.02E-11 | 3.02E-11 | 2.61E-10 |
| *nef* | | | 0.5997 | 3.65E-08 | 4.35E-05 | 1.16E-07 |

| | | | |
|---|---|---|---|
| *APOL1-G0* | 1.17E-09 | 4.69E-08 | 4.31E-08 |
| *APOL1-G0+nef* | | 0.001114 | 9.93E-02 |
| *APOL1-G1* | | | 7.20E-05 |
| *APOL1-G1+nef* | | | |
